# Conditionally Redundant Bacteriocin Targeting by *Pseudomonas syringae*

**DOI:** 10.1101/167593

**Authors:** Kevin L. Hockett, Meara Clark, Stacey Scott, David A. Baltrus

## Abstract

The widespread use of antimicrobials under clinical and agricultural settings has resulted in the evolution of resistance to these compounds. To combat the emergence of resistance, current research efforts are focusing on designing treatments to exploit combinations of antimicrobials, where the evolution of resistance confers sensitivity to alternative compounds. In this work we demonstrate that strains of *Pseudomonas syringae* possess a natural analogue to this strategy. Specifically, we demonstrate that a single strain produces multiple bacteriocins that can target another strain, but antimicrobial activity of the second bacteriocin is manifest only after resistance to the first emerges. The evolution of resistance also sensitizes the target strain to bacteriocins from a variety of other strains. Strains of *P. syringae* therefore encode multiple bacteriocins that can act in a conditionally redundant manner. It is possible that combinations of bacteriocins could be applied as a cocktail or sequentially, to potentially achieve durable pathogen control.

## Introduction

In many environments, bacteria reside in complex community assemblages, which can lead to intense competition for space and access to substrates^1–5^. To effectively compete for nutrients, bacteria frequently employ interference strategies, including the elaboration of diffusible anti-bacterial compounds, such as broad spectrum antibiotics ^6,7^ and bacteriocins^8–13^, or contact-dependent interference systems, such as the type IV, type V or type VI secretion systems^14–17^. Empirical and genomic data has shown that bacteria often encode multiple antagonistic traits, particularly among *Pseudomonas* strains^10,18–20^. In contrast to broad spectrum antibiotics, which have been widely used for nearly a century to combat and control bacterial pathogens under clinical and agricultural settings, bacteriocins have remained underutilized as pathogen control agents, despite documented potential^21–25^. This is partially because bacteriocins are thought to have a narrower spectrum of activity than more widely used antibiotics, which in the past has been considered detrimental but now is sought given worries about off-target effects on the microbiome. The notable exception to this overall trend being nisin, a small, posttranslationally modified peptide that has been incorporated into foods and food packaging as a spoilage preventative, as well as anti-*Listeria* compound^26–28^.

With the widely recognized breakdown in antibiotic-mediated control of human and agricultural pathogens^29–31^, resulting from the selection for and dissemination of antibiotic-resistance genes, there has been a push to both rethink how antibiotics are used, as well as the adoption of new control methods. To this end, researchers have explored several potentially intersecting approaches, none of which have yet been widely adopted, including: applying evolutionary principles to use antibiotics in temporal combinations to slow the emergence of resistance^32–34^, using antibiotic combinations that result in ‘collateral sensitivity’^35^, combining antibiotics with bacteriophages (phage-therapy) in a synergistic combination^36^, and utilizing bacteriocins (including combinations thereof) as narrow-spectrum antibacterials^21,23^. Inherent in all of these approaches is the recognition that, regardless of the selective agent, there are always paths to the evolution of resistance. Thus, there has been parallel interest in the development of strategies to counter or slow the evolution of antibiotic resistance and to harness trade-offs associated with resistance to bias microbial evolutionary trajectories towards those that involve a predictable fitness cost. As the vast majority of anti-microbials in our repertoire are produced by microbes, it is possible that microbes have already developed countermeasures to the challenge of competitors evolving antimicrobial resistance, and potentially evolved redundant control agents that exploit trade-offs associated with resistance.

One class of bacteriocins that has received recent attention are the bacteriophage-derived bacteriocins, termed tailocins^37^. These agents are evolutionarily derived from tailed bacteriophages. The best studied tailocins are the r-type and f-type pyocins of *Pseudomonas aeruginosa*^10,37,38^. The r-type tailocins are toxic toward target cells, exhibiting a one-hit-kill potency, where they cause depolarization of the target cell^39^. The r-type pyocins are divided into 5 subgroups based on their target strain tropism^40^, which is mediated by interactions between the tailocin tail fiber and the target cell lipopolysaccharide (LPS), the major constituent of Gram-negative outer membranes^41^. Domestication of phages into tailocins appears to be a common strategy employed by *Pseudomonas*, as there are 4 independently derived r-type tailocins across the genus^42,43^, with some strains producing two distinct r-type tailocins^44^. Importantly, tailocins can be modified to target important human pathogens, such as *E. coli* O157:H7, have been shown to be effective in reducing pathogen load within a mouse model^45,46^.

In this paper, we present evidence and argue that the ubiquitous plant-pathogen *Pseudomonas syringae* employs a conditionally redundant interference mechanism that exhibits the hallmarks of an evolutionarily robust control strategy. Specifically, gaining resistance to one inhibitory agent (an r-type syringacin or tailocin^43^) results in acquired sensitivity to an alternative s-type syringacin encoded by the same producing strain. To our knowledge, this is the first report of such conditional redundancy associated with naturally produced antimicrobials. Furthermore, we demonstrate that loss of LPS also sensitizes target strains to multiple types of killing activity by a variety of other *P. syringae* strains. These findings significantly inform our understanding of the evolution of antagonism within single bacterial species, and the mechanism of bacteriocin activity within Pseudomonads, but also suggest that forward thinking treatment regimes involving bacteriocins can be engineered as next generation pathogen control strategies.

## Results

P. syringae *pv*. actinidiae *BV3 exposed to supernatant of* P. syringae *pv*. syringae *B728a acquire tailocin resistance at significantly lower frequencies than other strains that are sensitive to the same tailocin*.

*P. syringae* pv. *syringae* B728a (*Psy*) produces a tailocin that targets several related strains of *P. syringae*, including *P. syringae* pv. *phaseolicola* 1448a (*Pph*), *P. glycinea* R4 (*Pgy*), *P. syringae* pv. *morsprunorum* (*Pmp*), and *P. syringae* pv. *actinidiae* J-1 (*Psa* J-1, synonym *Pan*)^43^. We found that an additional strain of *Psa, Psa* BV3, which is closely related to, but distinct from *Psa* J-1^47^, is also sensitive to the *Psy* tailocin (see below). In selecting for tailocin-resistant isolates, we found that *Pph, Pgy* and *Pmp* seemingly acquired resistance much more frequently than *Psa* BV3 (Fig. 1). This suggested that despite the tailocin being the sole source of killing present in the *Psy* B728a supernatant (tailocin deletion mutants do not inhibit *Psa* BV3 in an agar overlay assay, see below), that an additional toxic agent was inhibiting the emergence of tailocin resistant strains. In screening supernatants of *Psy* B728a derivative strains, which harbor deletions of predicted s-type bacteriocins, we found that resistant mutants readily emerged when exposed to the supernatant of *Psy* where an s-type bacteriocin (S_E9a_) has been deleted (Fig. 1). These results suggested that although *Psa* BV3 itself is not directly inhibited by S_E9a_, emergence of tailocin-resistant mutants might be inhibited by this alternative bacteriocin.

**Figure 1.**
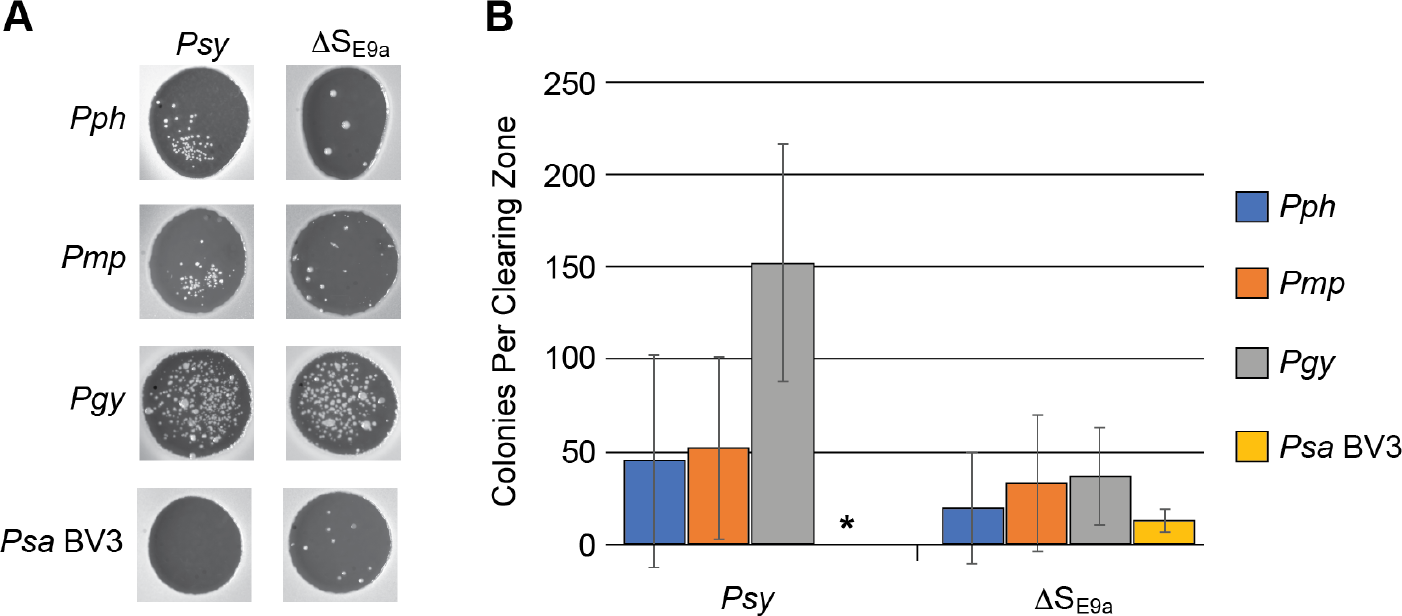
Emergence of bacteriocin resistant colonies of *Pph, Pmp, Pgy* or *Psa* BV3 following exposure to *Psy* supernatant. **A)** Representative killing zones with resistant colonies arising following 48 hours of incubation following exposure to *Psy* (containing both tailocin and s-type bacteriocins) or ΔS_E9a_ (containing tailocin, but lacking the S_E9a_ s-type bacteriocin). **B)** Quantification of the number of resistant colonies arising from exposure either to the *Psy* or ΔS_E9a_ supernatants. Error bars indicate standard of the mean across six replicate zones. *No colonies were observed in the *Psy* killing zones on *Psa* BV3 overlays. This experiment was repeated twice with similar results.

Psa *BV3 strains with transposon insertions in LPS biosynthetic genes exhibit a loss of O-antigen bands and shifted bacteriocin sensitivity*.

Previous work has demonstrated that tailocin-linked killing is widespread across *Pseudomonas syringae*^43^. Although evolutionarily distinct from the *P. syringae* tailocin, an analogous tailocin encoded by *P. aeruginosa* (r-type pyocin) has been reported use either lipopolysaccharide (LPS) or lipooligosaccharide (LOS) as the cell surface receptor^41,48^. To determine whether LPS serves as the cell surface receptor for the *P. syringae* tailocin, we screened a collection of strains harboring transposon insertions in genes involved in LPS biosynthesis^49^ for resistance to the *Psy* B728a tailocin (Fig. 2). Although the wild type strain (BV3), strains with altered LPS biosynthesis, and strains with transposon insertions in genes unrelated to LPS biosynthesis were all inhibited by *Psy* culture supernatant, the inhibitory phenotype of the LPS mutants was markedly different than that for the strains with unaltered LPS (Fig. 2). The clearing zones for BV3, A12, and C2 (see Table 1 for strain genotypes) treated with *Psy* supernatant exhibit a crisp, defined edge, which is characteristic of tailocin-mediated killing. We confirmed that the LPS isolated from mutants A12 and C2 were similar to the wild-type BV3 strain (Fig. 3). For all strains with altered LPS, however, clearing zones resulting from the same supernatant had a diffuse, ill-defined edge. These results indicated the clearing for the latter strains did not result from tailocin-mediated killing, but likely from an alternate, low-molecular weight bacteriocin^43,50^. All of the strains we tested that harbor transposon insertions in predicted LPS biosynthetic genes exhibited loss of their O-antigen component (Fig. 3). To confirm the diffuse clearing zones did not result from tailocin-mediated killing, we tested for killing activity from strains that do not produce active tailocins, ΔR_struct_ and ΔR_rbp_^43^. These strains do not kill *Psa* BV3, A12 or C2, but do retain killing activity against LPS mutants. *Psy* supernatant that has undergone polyethylene glycol (PEG) precipitation and resuspension, which efficiently recovers large protein complexes, but not smaller, single proteins, retains killing activity against wild-type LPS strains only. Conversely, deletion of S_E9a_, encoding a single protein, s-type bacteriocin with a predicted DNase catalytic domain^43,50^ results in loss of killing activity against the LPS mutants only. Taken together, these results clearly demonstrate that *Psy* produces two distinct bacteriocins that are conditionally redundant in targeting *Psa* BV3.

**Figure 2.**
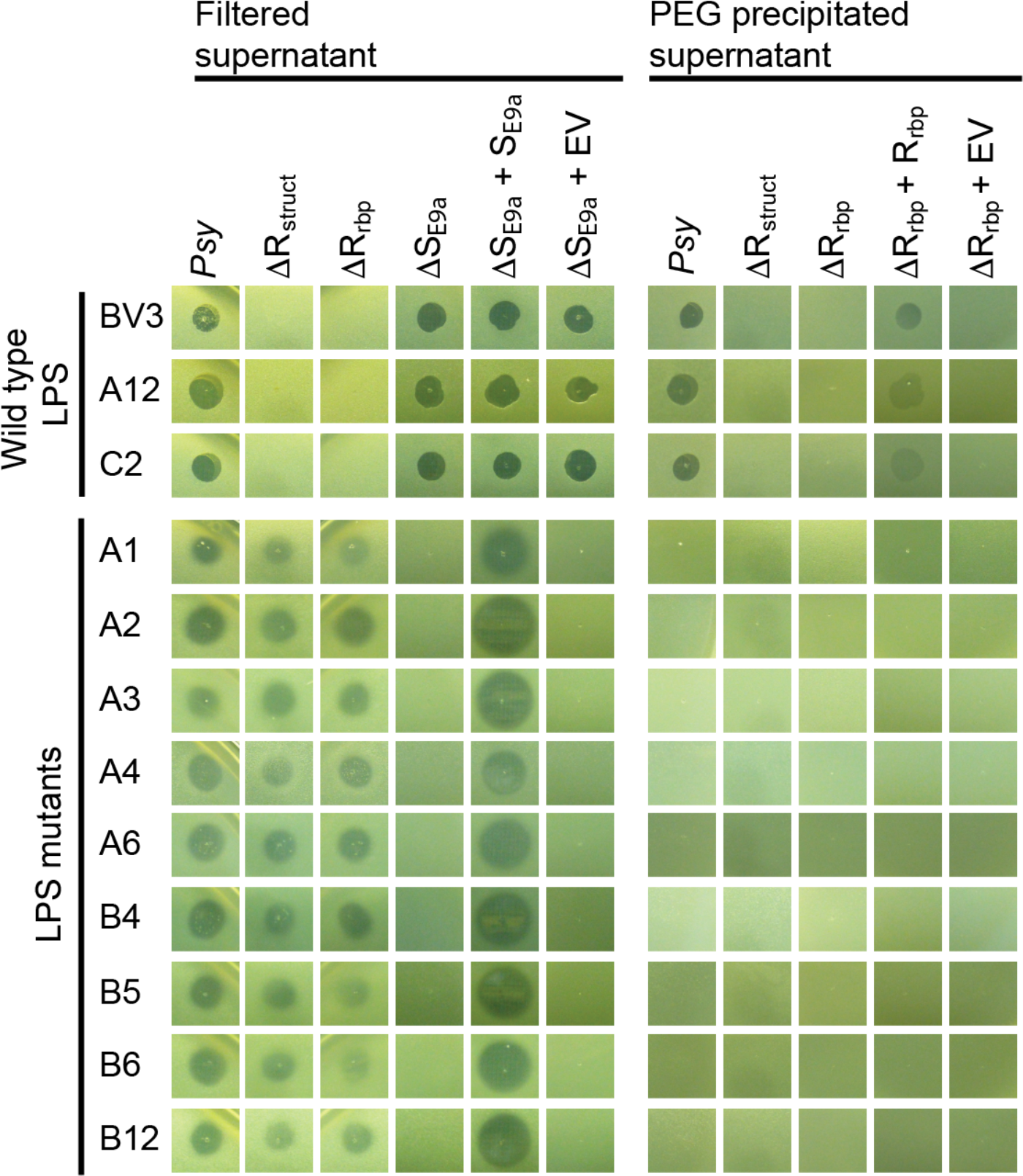
*Psa* strains harboring LPS mutations acquire sensitivity to an alternative, s-type syringacin produced by *Psy* B728a. Row labels indicate *Psa* overlay strain (see Table 1 for strain genotypes) and whether the LPS is altered or not (see Fig. 3). Column labels indicate the genotype of the supernatant source (*Psy*, wild type *Psy* B728a) and whether the supernatant was partially purified by PEG precipitation (see methods) or not. See Table 1 for *Psy* B728a genotypes.

**Figure 3.**
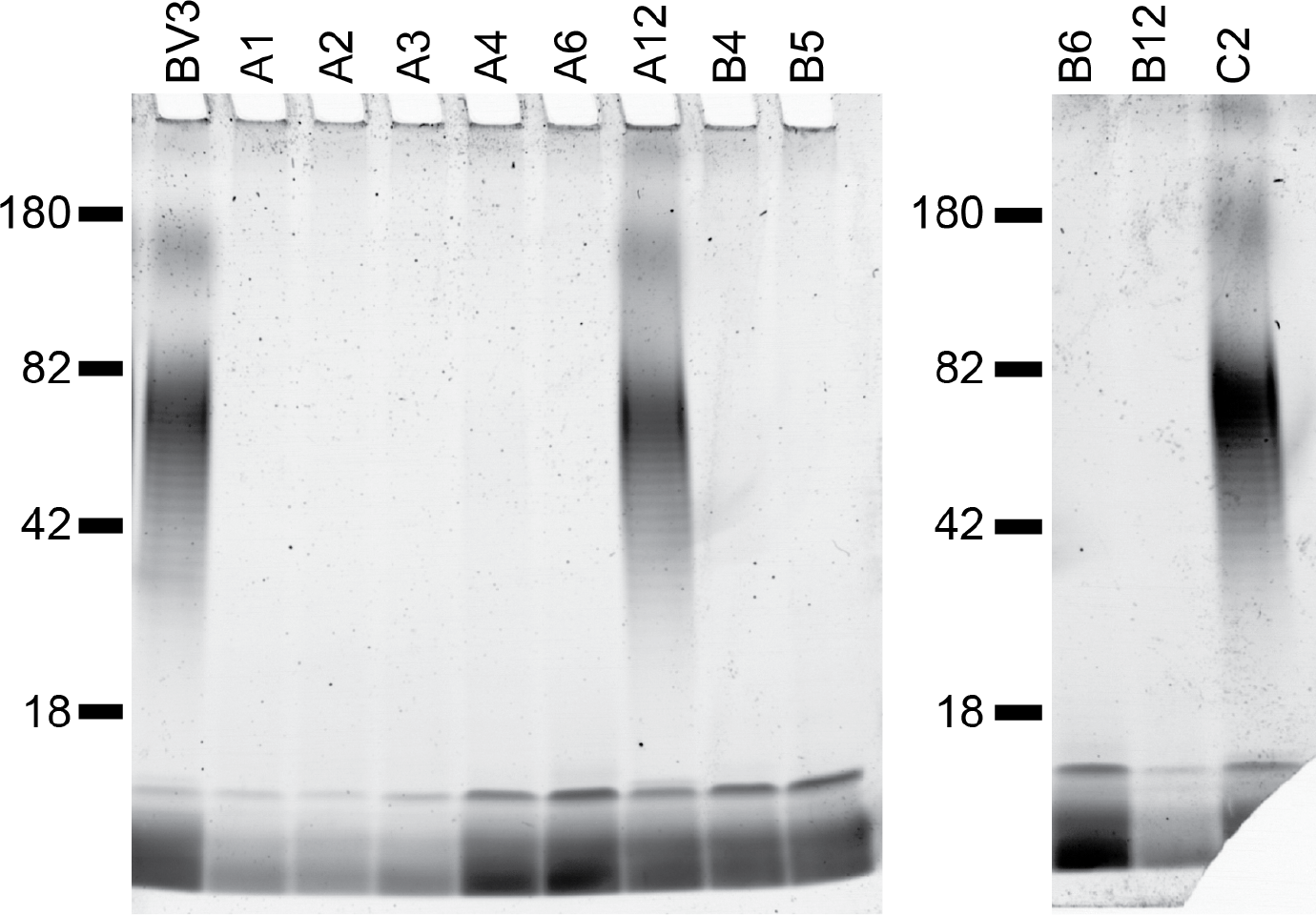
Strains exhibiting acquired sensitivity to the s-type syringacin of *Psy* B728a have lost O-antigen bands. Isolated LPS preparations were separated by SDS PAGE and stained using Pro-Q Emerald 300. Marker sizes (kDa) are indicated.

**Table 1.** Strains and Plasmids

| Strain or plasmid | Description or Relevant Characteristics | Antibiotics | Reference |
| --- | --- | --- | --- |
| *E. coli* strains |  |  |  |
| TOP10 | F- *mcrA* $\Delta$(*mrr-hsdRMS-mcrBC*) ϕ80*lacZ*ΔM15 Δ*lacX74* *recA1* *araD139* $\Delta$(*araleu*) 7697 *galU* *galK* *rpsL* (Str<sup>R</sup>) *endA1* *nupG* | | Invitrogen |
| DAB44 | *E. coli* harboring pRK2013, (conjugation helper for triparental mating) |  | 79 |
| *P. syringae* strains |  |  |  |
| Psy B728a | *P. syringae* pv. *syringae* B728a | Rif<sup>R</sup> | 50 |
| ΔS<sub>E9a</sub> | Psy B728a harboring a markerless deletion of Psyr_4651 and Psyr_4652 (previously labelled Δ4651) | Rif<sup>R</sup> | 43 |
| ΔS<sub>E9a</sub> + pS<sub>E9a</sub> | $\Delta$ S<sub>E9a</sub> harboring pS<sub>E9a</sub> complementation vector | Rif<sup>R</sup>, Kan<sup>R</sup> | This work |
| ΔS<sub>E9a</sub> + pJC531EV | $\Delta$ S<sub>E9a</sub> harboring pJC531EV | Rif<sup>R</sup>, Kan<sup>R</sup> | This work |
| ΔR<sub>struct</sub> | Psy B728a harboring a markerless deletion of Psyr_4586 through Psyr_4592 and the promoter of Psyr_4585 | Rif<sup>R</sup> | 43 |
| ΔR<sub>rbp</sub> | Psy B278a harboring a markerless deletion of Psyr_4585 and Psyr_4584 | Rif<sup>R</sup> | 43 |
| ΔR<sub>rbp</sub> + pR<sub>rbp</sub> | ΔR<sub>rbp</sub> harboring pR<sub>rbp</sub> complementation vector | Rif<sup>R</sup>, Tet<sup>R</sup> | 43 |
| ΔR<sub>rbp</sub> + pBAV226EV | ΔR<sub>rbp</sub> harboring pBAV226EV | Rif<sup>R</sup>, Tet<sup>R</sup> | This work |
| Psa BV3 | *Pseudomonas syringae* pv. *actinidiae* New Zealand isolate PsaV (ICMP 18884; Biovar 3) |  | 80 |
| A1 | Psa BV3; Tn insertion into IYO_023030 (glycosyl transferase family 1); LPS mutant | Kan<sup>R</sup> | 80 |
| A2 | Psa BV3; Tn insertion into IYO_025555 (O-antigen ligase, Wzy); LPS mutant | Kan<sup>R</sup> | 80 |
| A3 | Psa BV3; Tn insertion into IYO_008995 (glycosyl transferase); LPS mutant | Kan<sup>R</sup> | 80 |
| A4 | Psa BV3; Tn insertion into IYO_023015 (O-antigen export system permease protein RfbD); LPS mutant | Kan<sup>R</sup> | 80 |
| A6 | Psa BV3; Tn insertion into IYO_023005 (SAM-dependent methyltransferase/WbbD); LPS mutant | Kan<sup>R</sup> | 80 |
| A12 | *Psa* BV3; Tn insertion into IYO_018250 (glycerol-3-phosphate dehydrogenase); non-LPS mutant | Kan<sup>R</sup> | <sup>80</sup> |
| B4 | *Psa* BV3; Tn insertion into IYO_023010 (sugar ABC transporter); LPS mutant | Kan<sup>R</sup> | <sup>80</sup> |
| B5 | *Psa* BV3; Tn insertion into IYO_005330 (glycosyl transferase); LPS mutant | Kan<sup>R</sup> | <sup>80</sup> |
| B6 | *Psa* BV3; Tn insertion into IYO_023005 (glycosyl transferase); LPS mutant | Kan<sup>R</sup> | <sup>80</sup> |
| B12 | *Psa* BV3; Tn insertion into IYO_023005 (glycosyl transferase); LPS mutant | Kan<sup>R</sup> | <sup>80</sup> |
| C2 | *Psa* BV3; Tn insertion into IYO_009915 (flagellar p-ring protein flgl); non-LPS mutant | Kan<sup>R</sup> | <sup>80</sup> |
| *Pph* | *Pseudomonas syringae* pv. *phaseolicola* 1448a |  | <sup>81</sup> |
| *Pgy* | *Pseudomonas syringae* pv. *glycinea* R4 |  | <sup>81</sup> |
| *Pmp* | *Pseudomonas syringae* pv. *morsprunorum* |  | <sup>81</sup> |
| USA011 | *Pseudomonas syringae* USA011 |  | <sup>82</sup> |
| CC1416 | *Pseudomonas syringae* CC1416 |  | <sup>82</sup> |
| CC1466 | *Pseudomonas syringae* CC1466 |  | <sup>82</sup> |
| UB303 | *Pseudomonas syringae* UB303 |  | <sup>82</sup> |
| CC94 | *Pseudomonas syringae* CC94 |  | <sup>83</sup> |
| TLP2 | *Pseudomonas syringae* TLP2 |  | <sup>84</sup> |
| *Por* | *Pseudomonas syringae* pv. *oryzae* 1_0 |  | <sup>81</sup> |
| *Pae* | *Pseudomonas syringae* pv. *aesculi* |  | <sup>81</sup> |
| CC1543 | *Pseudomonas syringae* CC1543 |  | <sup>82</sup> |
| *Pja* | *Pseudomonas syringae* pv. *japonica* |  | <sup>81</sup> |
| Plasmids |  |  |  |
| pJC531 | Replicative destination vector harboring *attR* cloning sites | Kan<sup>R</sup> | <sup>85</sup> |
| pDONR207 | Entry vector harboring *attP* cloning sites | Gent<sup>R</sup>,<br>Cam<sup>R</sup> | Invitrogen |
| pBAV226 | Replicative destination vector with constitutive nptII promoter | Tet<sup>R</sup> | <sup>86</sup> |
| pR<sub>rbp</sub> | Psyr_4584 and Psyr_4585 complementation construct (including upstream alternative start codon) in pBAV226 |  | <sup>43</sup> |
| pS<sub>E9a</sub> | Psyr_4651 and Psyr_4652 complementation construct (including upstream promoter region) cloned into pJC531 | Kan<sup>R</sup> | This work |
| pJC531EV | pJC531 with a short stretch of non-coding DNA integrated, serving as an empty vector control | Kan<sup>R</sup> | <sup>85</sup> |
| pBAV226EV | pBAV226 with a short stretch of non-coding DNA integrated, serving as an empty vector control | Tet<sup>R</sup> | <sup>86</sup> |

Since LPS mutants of *Psa* BV3 were sensitized to alternative bacteriocins from *Psy*, we tested whether a LPS mutants were also sensitized to bacteriocins produced by variety of other *P. syringae* strains. We first screened wild-type BV3 against a diverse collection of *P. syringae* species complex strains isolated from a variety of sources (Figure 4 and Supplemental Table 1) using the same overlay assays described above, and found that numerous strains produce killing activity that is consistent with tailocins. This activity creates crisp edges in the overlay assay, is retained following PEG precipitation, and is eliminated in the LPS mutants. In a small number of cases it appeared that strains contained s-type bacteriocins that could target BV3 (i.e. from strain UB303) and this activity remained in the LPS mutants. We also found a case (CC1543) where there was no killing activity in the wild type strain but where an s-type activity was uncovered in the LPS mutants. Lastly, and surprisingly, we found numerous cases where a tailocin-like activity (with crisp edges and precipitated by PEG) was uncovered against the LPS mutants but not the wild type strains. This last data point strongly suggests that, while many and perhaps a majority of r-type syringacins target LPS, a subset can target an alternative cell surface receptor(s) that are inaccessible in a wild-type LPS background.

**Figure 4.**
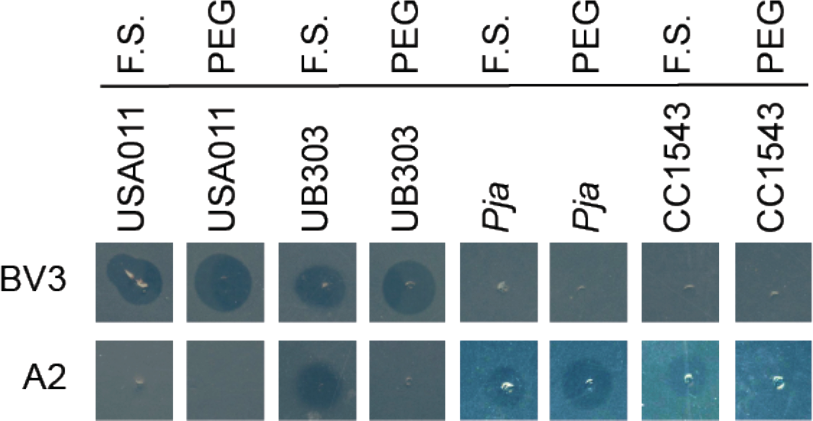
Differential bacteriocin-mediated interactions between *P. syringae* strains and either wild-type (BV3) or an LPS mutant (A2) of *Psa*. Filter sterilized supernatants were applied directly (F.S.) or PEG precipitated (PEG). Putatively, these results indicate: USA011 produces an O-antigen-targeting tailocin only; UB303 produces an O-antigen-targeting tailocin and an s-type bacteriocin that targets an O-antigen null mutant; *Pja* produces a tailocin that targets an O-antigen null mutatnt; CC1543 produces an s-type bacteriocin that targets an O-antigen null mutant.

## DISCUSSION

Over the past several decades, the rise in antibiotic resistance has necessitated the development of alternative pathogen control strategies for both medical and agricultural settings. These next generation antimicrobial treatments include the adoption of antibiotic alternatives, such as bacteriocins, lytic bacteriophages or CRISPR-Cas systems^51,52^, rationally designed antibiotic regimens, and a combination of antibiotic and non-antibiotic agents. The overarching goal of each approach, however, focuses on how best to slow the emergence of resistance and control breakdown. Bacteriophages have been employed successfully in certain instances to treat animal^53–55^ and plant^56^ infections, and are thought to be particularly advantageous over traditional antibiotics because of their ability to amplify in the presence of their target host^57^ and to overcome resistance through reciprocal coevolution. However, concerns remain, regarding the ability of phages ability to both promote horizontal gene transfer of potentially pathogenic determinants and to shift host tropism^58^. Alternatively, researchers have begun to explore the development of combined antimicrobial treatment regimes whereby resistance to one compound leads to sensitivity to a second antibacterial. This strategy is limited by the recognition that the use of multiple, broad spectrum antimicrobials can eliminate beneficial members of the microbiome and render hosts susceptible to secondary infections^59^. Recently, a strategy has been successfully employed that combines both of these ideas as the deployment of phages that use antibiotic efflux pumps as a cell surface receptor has been shown to result bacterial strains that trade phage resistance for antibiotic sensitivity^60^.

In this paper, we describe a strain of *P. syringae* that naturally encodes two distinct bacteriocins (an r-type and s-type syringacin) that redundantly target the same strain. However, this redundancy is conditional, in that the s-type syringacin is only effective against strains that are resistant to the r-type syringacin. Intriguingly, these results suggest this strain has arrived at an evolutionary result broadly in line with theoretical and empirical research focusing on robust control strategies. This strain is predicted to encode two additional s-type syringacins for which, to our knowledge, no target strains have been found but which could factor into further layers of redundancy. Given the widespread occurrence of tailocin-mediated killing and diffusible bacteriocins encoded by Pseudomonad genomes, redundant targeting may exist broadly across this species, if not genus^10,18,43^. Indeed, in screening a limited but phylogenetically diverse set of *P. syringae* species complex strains we have found multiple instances where LPS mutants of *Psa* BV3, but not the wild type, are sensitized to bacteriocin-like killing from other strains. Sensitization appears to occur to s-type bacteriocins and, in one case (supernatant from *P. syringae* pv. *japonica*) to killing activity that is consistent with r-type tailocins. If this latter activity is due to an r-type tailocin, it also implies that there may be alternative targeting sites for these bacteriocins other than the LPS. Furthermore, this conditionally redundant phenomenon was briefly noted in a review on bacteriophage-and bacteriocin-mediated control of phytopathogens^61^, though these results appear to have never made their way into the primary literature. Vidaver noted the majority of isolates gaining resistance to syringacin 4-A (a tailocin^62^) acquired sensitivity to syringacin W-1 (also a tailocin^63^).

From the viewpoints of both evolutionary biology and classical microbial genetics, it is interesting to note that there was a striking difference in emergence of tailocin resistance in *Psa* BV3 when exposed to the tailocin alone or in combination with S_E9a_. Such differences in rates of mutation can be attributed to sensitization as a cost of resistance, but they may not be the only costs manifest under natural conditions. In other research on tailocin resistance in *P. syringae*, we have found that tailocin resistant mutants also exhibit a pronounced reduction in proliferation within their host plants (Hockett and Baltrus, in prep). Whether a similar reduction in virulence occurs for tailocin-resistant *Psa* BV3 when infecting its host plant, kiwifruit, remains untested. Taken together, our results suggest a cocktail of the r-type and s-type syringacin could be a potent control strategy for *Psa* BV3, which has resulted in significant loss of kiwi production, and associated economic loss, in New Zealand^64^. This control strategy would be all the more effective, if bacteriocin resistance resulted in a pleiotropic loss of fitness within the plant environment, which we hypothesize will occur.

Furthermore, we have demonstrated that other *P. syringae* species complex strains produce bacteriocins that can target LPS mutants of strain BV3 but not the wild type. This result has multiple important implications in terms of bacteriocin typing as well as biocontrol using bacteriocins. A better understanding of how context determines bacteriocin sensitivity, especially if there are multiple targets for tailocins, could enable the development of “smart” bacteriocin treatment strategies or the engineering of prophylactic bacteria with redundant killing capabilities.

Conditionally redundant inhibition may not rely strictly on genetic resistance. Although LPS is well described for its role in protecting against generally inhibitory substances, such as small antimicrobial peptides, and antibiotics, it has recently been shown to play role in protecting against an s-type bacteriocin in *E. coli*^65^. Growth within the biofilm environment both promotes the production of this bacteriocin, as well as stimulates the reduction in O-antigen length of the target strain, making this strain more sensitive. Thus, the growth conditions, and their effect on the physiology of both the producing and target strain, are likely to be important determinants of bacteriocin activity under environmentally relevant conditions. Although the cell surface receptor is currently unknown for the s-type syringacin, we have shown that LPS is likely the receptor for the tailocin. For other s-type bacteriocins encoded across *Pseudomonas*, either siderophore importers^66–69^ or LPS^70^ serves as the bacteriocin receptors. Growth conditions, specifically iron availability, can influence the sensitivity to these bacteriocins^71^. This might indicate there are growth conditions where the s-type syringacin is operative, even in the absence of genetic r-type syringacin resistance. Additionally, factors such as growth phase, transient alterations in gene expression, and nutrient availability are all known to affect either LPS biosynthesis^72–74^ and could potentially influence trade-offs in bacteriocin sensitivity.

## MATERIAL AND METHODS

### Plasmids, primers, bacterial isolates, and growth conditions

Bacterial strains and plasmids used in this study are listed in Table 1. All *P. syringae* strains were routinely grown on King’s medium B at 27 °C ^75^. *Eschericia coli* was routinely grown in lysogeny broth (10g NaCl/L)^76^ at 37 °C. When appropriate, growth media was amended with the antibiotics at the following concentrations: tetracycline (10 μg/ml), kanamycin (50 μg/ml), gentamicin (25 μg/ml), rifampin (50 μg/ml).

### Bacteriocin isolation and PEG precipitation

Bacteriocin samples were isolated as described ^43,77^. Briefly, overnight KB broth cultures of *Psy* B728a and derived mutants were diluted 1/100 into fresh KB medium and allowed to incubate for 2-4 hrs with shaking, after which mitomycin C was amended to the culture (0.5 μg/ml final concentration). The culture was incubated for an additional 24 hrs with shaking. Following incubation, the cell debris and surviving cells were removed by centrifugation (20,000 x g for 5 min) and filter sterilization (0.22 μm pore size) of the supernatant. We employed a polyethylene glycol (PEG) precipitation step for certain sterilized supernatants to preferentially recover high molecular weight bacteriocins (i.e. tailocins). To do this, PEG 8000 and NaCl were added at 10% (w/v) and 1 M final concentrations, respectively, then allowed to incubate in an ice bath for 1 hr. Samples were then centrifuged at 16,000 x g at 4 °C for 30 min. After decanting and discarding the supernatant, the pellet was resuspended into 1/10 the original volume in buffer (10 mM Tris, 10 mM MgSO4, pH 7.0). Residual PEG was removed by sequential chloroform extractions until no white interface was visible between the organic and aqueous phase.

### Agar overlay assay

The agar overlay assay was performed as previously described ^43,77^. Briefly, strains to be tested for bacteriocin sensitivity were cultured overnight in KB broth with shaking. The following morning the culture was diluted 1/100 into fresh KB broth and allowed to incubate with shaking for 2-4 hours. Following incubation, 100 μl of each culture was inoculated separately into 3 ml of molten 0.35% water agar. The bacterial suspensions were spread evenly over separate KB agar plates and allowed to solidify covered at room temperature for 30 min. To each solidified plate, 2-5 μl of bacteriocin preparation was spotted. Overlay plates were allowed to incubate overnight on the bench top, with results were recorded the following day.

### LPS extraction and visualization

LPS was extracted and visualized as described ^78^. Briefly, cultures were grown overnight in KB broth with shaking. The following morning, strains were diluted 1/100 into fresh KB broth and incubated for 2-4 hours. Following this incubation, 1 ml of culture was pelleted in a 1.5 ml microfuge tube (10,600 x g for 10 min.) and the supernatant was discarded. The pellet was resuspended in 200 μl 1x SDS buffer (2% *β*-mercaptoethanol [BME], 2% SDS and 10% glycerol in 50 mM Tris-HCl, pH 6.8, with a pinch of bromophenol blue dye) by repeated pipetting. The suspension was boiled for 15 min. Then allowed to cool to room temperature. 5 μl of DNase I and RNase solutions (both 10 mg/ml) were added to the boiled suspension and incubated for 30 min at 37 °C. 10 μl of a Proteinase K solution (10 mg/ml) was added to each sample, followed by incubation at 59 °C for 3 hrs. Samples were then extracted by combining with 200 μl of ice-cold Tris-saturated phenol, vortexing for 10 sec, then incubating at 65 °C for 15 min. Following this incubation, samples were cooled to room temperature and 1 ml of diethyl ether was added and vortexed for 10 seconds. The sample was then centrifuged at 20,600 x g for 10 min. The bottom aqueous layer (blue in coloration because of the bromophenol blue dye) was removed and extracted one additional time with chloroform and diethyl ether. After the final extraction, 200 μl of 2x SDS buffer was added and the samples were separated by SDS-PAGE. Samples were labeled using the Pro-Q Emerald 300 glycoprotein stain kit (ThermoFisher Scientific, #P21857) per the manufacturer’s instructions. Samples were visualized using Azure c600 gel imaging system. CandyCane™ glycoprotein molecular weight standards (ThermoFisher Scientific, #C21852) was used for LPS size determination.

## ACKNOWLEDGMENTS

We thank Matt Templeton for kindly sharing Psa BV3 and LPS mutant derivative strains. This research was supported by the National Intsitute of Food and Agriculture, U.S. Department of Agriculture, under award number 2016-67014-24805.

## AUTHOR CONTRIBUTIONS

K.L.H. and D.A.B. designed research; K.L.H., M.C. and S.S. performed the research; K.L.H. and D.A.B. analysed data and wrote the manuscript.

## CONFLICTS OF INTEREST

Authors declare no conflict of interest.

## SUPPLEMENTARY TABLES

**Supplementary Table 1.** Effect of LPS mutation on bacteriocin mediated killing by diverse *P. syringae* strains.

|  | WT *Psa* BV3 |  | LPS mutants |  |
| --- | --- | --- | --- | --- |
|  | Filtered Supernatant | PEG Precipitated | Filtered Supernatant | PEG Precipitated |
| *Psy* B728a | + | + | + | - |
| USA011 | + | + | - | - |
| CC1416 | + | + | - | - |
| CC1466 | + | + | - | - |
| UB303 | + | + | + | - |
| CC94 | + | + | - | - |
| TLP2 | + | + | - | - |
| *Por* | + | + | - | - |
| *Pae* | - | (+) | - | - |
| CC1543 | - | - | + | - |
| *Pja* | - | - | + | + |

+ clearing zone; - no clearing zone; (+) faint clearing zone

